# Beyond tortoises and hares: Niche lability and a continuum of diversification dynamics underlie the megadiversity of a legume genus

**DOI:** 10.64898/2026.11.06.764677

**Authors:** Rafael F. Barduzzi, Yago Barros-Souza, Daniel S. Moen, Leonardo M. Borges

## Abstract

- Variation in macroevolutionary dynamics, from rapid radiations to slower evolutionary change, can shape multiple facets of biodiversity. However, whether the contributions of contrasting macroevolutionary dynamics to biodiversity differ across timescales, and whether shifts in abiotic niches drive recent radiations, remain open questions in evolutionary biology.
- Using the megadiverse genus *Mimosa*, we investigate the contribution of contrasting macroevolutionary dynamics to biodiversity and what drives these dynamics at shallow timescales. We combined species-diversity, morphological, and abiotic niche data to estimate biodiversity metrics, infer macroevolutionary rates, and test the role of abiotic niche lability as a driver of radiation-like dynamics.
- We found that *Mimosa* clades are distributed across a continuum of macroevolutionary dynamics, with those clades resembling radiations clearly contributing to most diversity and disparity, but not niche breadth. Moreover, rates of altitudinal and edaphic niche change were associated with diversification, whereas rates of change in these two niche dimensions, along with climatic niche change, were associated with morphological evolution. Finally, the contribution of radiations to biodiversity seemed to decrease toward the present.
- Overall, our work highlights that both relatively rapid and slow lineage diversification underlie the multifaceted nature of biodiversity and reinforces the role of abiotic niche lability in plant diversification.

## Introduction

Our understanding of tempo in macroevolution, from phenotypic change to lineage formation, has advanced greatly through studies examining the contrast between rapid and slow dynamics (Simpson, 1953; Erwin, 1992; Olson & Arroyo-Santos, 2009). For instance, the study of clades exhibiting rapid morphological evolution or lineage accumulation has uncovered some of the drivers of accelerated biological diversification across the tree of life (Simões *et al*., 2016). Such events appear to be triggered and maintained by a multitude of intrinsic and extrinsic factors, such as ‘key’ changes in phenotype, ecology, or the landscape where species occur (Givnish, 2015; Stroud & Losos, 2016; Simões *et al*., 2016). However, clades with significantly slower diversification dynamics (e.g. lineage ‘depauperons’ *sensu* Donoghue & Sanderson, 2015) are also common across the tree of life, and understanding the processes shaping this pattern may provide insights into the constraints on biological diversification (Donoghue & Sanderson, 2015). For example, do such contrasting dynamics (i.e., clades that diversify faster or slower over time) contribute differentially to multiple facets of biodiversity, such as species diversity, morphological diversity (hereafter ‘disparity’), and niche breadth (Whittaker, 1972; Wills *et al*., 1994)?

Early ideas on macroevolutionary dynamics suggested that clades displaying rapid lineage diversification, commonly referred to as ‘radiations’, represented an extreme of the continuum that contributed to most of the diversity of life (Simpson, 1953; Glor, 2010; Donoghue & Sanderson, 2015; Hughes *et al*., 2015). The few studies that have explicitly addressed this prediction found that rapid diversification indeed underlies most species diversity and disparity (of ecologically relevant traits) over deep time, from 1.5 billion years ago (Ga) to 80 million years ago (Ma), encompassing taxonomic levels ranging from kingdoms to families (Morinaga *et al*., 2023; Wiens & Moen, 2025). Nonetheless, the relative contributions of different temporal dynamics of diversification to the generation of biodiversity may differ among taxonomic groups and evolutionary timescales (Hiller *et al*., 2019; Stephens *et al*., 2025). Such differences may reflect variation in intrinsic and extrinsic factors affecting diversification and evolutionary trajectories (Jablonski, 2017), as well as differences in the biodiversity metrics being evaluated and the ways in which they are measured. Indeed, for squamates, the correlation between diversification rates and species richness weakens in time frames nearest to the present—that is, within the last ∼20 Ma (Stephens *et al*., 2025). However, the role of different macroevolutionary dynamics in generating biodiversity (including species diversity, disparity, and niche breadth) at these shallower timescales remains to be tested for most groups of organisms.

A promising framework for addressing the relationship between macroevolutionary rate dynamics and the generation of biodiversity lies in the inference of ‘radiation spaces’. This approach investigates how clades are distributed across a space defined by a spectrum of variation in macroevolutionary rates, such as rates of lineage diversification and phenotypic evolution (Sanderson, 1998; Lieberman, 2012; Moen *et al*., 2021; Morinaga *et al*., 2023). Based on their position in pre-defined areas of the radiation-space continuum (e.g. quadrants in a bidimensional space), lineages are then hypothesized to represent distinct macroevolutionary dynamics, such as adaptive radiations (high rates of both lineage and phenotypic evolution) or non-adaptive radiations (high rates of lineage diversification, but not phenotypic evolution). Additionally, estimating biodiversity metrics (e.g. species diversity and morphological disparity) for each area of the radiation space summarizes the contribution of each type of macroevolutionary dynamic to the generation of current biodiversity. So far, studies that applied such a framework to broad taxonomic groups and deep evolutionary timescales found that radiations account for much of extant species diversity and eco-morphological disparity. This approach is therefore well suited to examining whether the importance of radiations in generating biodiversity extends to shallower evolutionary timescales and additional dimensions of biodiversity, such as niche breadth.

Niche evolution is expected to influence diversification dynamics and is at the core of hypotheses aiming to explain variation in diversity among groups (Simpson, 1953; Stroud and Losos, 2016). Indeed, examples from both plants (e.g. Andean Lupinus and Hawaiian lobeliads; Hughes & Eastwood, 2006; Givnish *et al*., 2009; Drummond et al., 2012) and animals (e.g. cichlid fishes and Malagasy vangas; Seehausen, 2015; Ford *et al*., 2016; Auerbach *et al*., 2025) invoke habitat shifts as some of the main drivers of radiations, sometimes acting together with morphological innovations (Liem, 1973; Nürk *et al*., 2019). However, despite growing interest in how these intertwined processes shape plant diversification, studies explicitly examining the relationships between the rates of diversification, phenotypic evolution, and abiotic niche change remain relatively scarce (Folk *et al*., 2019; Qi *et al*., 2023; Dellinger *et al*., 2024). Moreover, simulations aiming to understand how biogeography drives radiations suggest that niche partitioning may be a consequence of such dynamics (Tao *et al*., 2021). Given this context, integrating environmental variables into analyses of macroevolutionary dynamics provides an opportunity to better understand the relationship between niche diversity and lineage diversification, including the extent to which niche diversity may act as a driver of diversification.

Highly diverse plant genera provide valuable systems for investigating the drivers and outcomes of contrasting macroevolutionary dynamics at shallower evolutionary timescales. Their often recent diversification histories (∼30–20 Ma), combined with high species diversity and frequently substantial morphological and niche diversity, allow studies of how lineage diversification, phenotypic evolution, and niche changes jointly generate biodiversity (Frodin, 2004; Dimitrov et al., 2023; Moonlight et al., 2024). For instance, *Lupinus* L. and *Euphorbia* L. show multiple diversification dynamics, including well-known cases classified as evolutionary radiations. In these cases, the combination of phenotypic changes (e.g. in morphology or physiology) and colonization of new abiotic niches (e.g. along altitudinal or climatic gradients) has been hypothesized to underlie radiations (Drummond et al., 2012; Horn et al., 2014). Such examples align well with the currently most widely supported hypotheses concerning the drivers of plant evolution, which posits that rapid diversification may arise either from the combined effect of sequentially evolved traits (’key synnovation’) or from the interaction between such traits and colonization of new environments (’key confluence’; Donoghue & Sanderson, 2015; Cai *et al*., 2025).

Among legumes, the third-largest family of flowering plants (LPWG, 2017), *Mimosa* L. is a genus that stands out for its taxonomic, morphological, and ecological diversity (Fig. 1). The genus includes 600+ species, most of which occur across the tropical and subtropical Americas, while 32 species are endemic to Madagascar and 8 occur in Asia and continental Africa (Barneby, 1991; Villiers, 2002; Borges *et al*., 2024). *Mimosa* has high vegetative diversity, including trees, shrubs, and climbers with ample variation in the subdivision of their bipinnately compound leaves (Fig. 1). Moreover, *Mimosa* occurs in a wide array of biomes, including tropical forests, savannas, deserts, grasslands, and wetlands (Fig. 1; Borges *et al*., 2024; Simon *et al*., 2011). Finally, the evolutionary history of the genus includes multiple niche shifts (e.g. transitions between tropical forest and savanna) and diversification bursts, such as those in lineages from Central Brazil and Madagascar (Barneby, 1991; Koenen *et al*., 2013; Vasconcelos *et al*., 2020; Borges *et al*., 2023; Ringelberg *et al*., 2023; Cássia-Silva *et al*., 2026). *Mimosa* thus stands out as a promising group for studying both the contributions of radiations to plant diversity and their drivers.

**Fig. 1.**
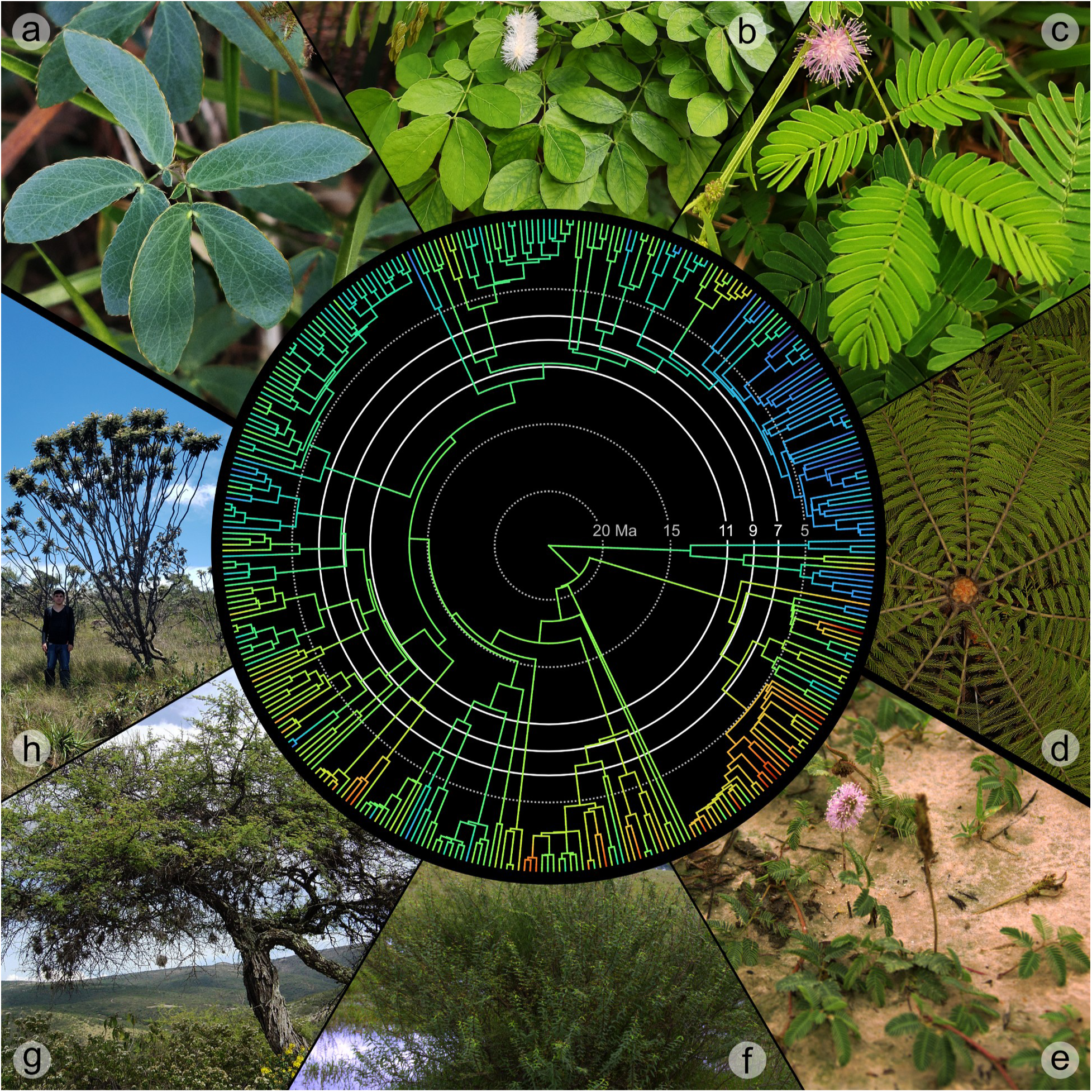
Facets of *Mimosa* biological diversity. The branches of the genus-level phylogeny show an ancestral-state estimation for PC1 of our dataset of abiotic variables. The rainbow scale varies from low PC1 values (red and yellow; warmer conditions with sandier, more alkaline, and denser soils) to high PC1 values (green and blue; wetter conditions with higher soil organic carbon and nitrogen). The ages used to delimit clades in this study (11, 9, and 7 million years ago) are indicated among the temporal markers. Photographs around the edges highlight variation in both vegetative morphology and niche across the genus. (a-d) Leaf morphological diversity, with varying degrees of leaf subdivision. (e-h) Habit and ecological diversity, from prostrate subshrubs to trees, occupying environments characterized by sandy soils, wetlands, grasslands, and savannas. The photographs are arranged to illustrate gradients of morphological change rather than phylogenetic relationships; therefore, their positions in the figure do not correspond to the positions of the respective taxa in the phylogeny. (a) *Mimosa debilis*, (b) *M. caesalpiniifolia*, (c) *M. candollei*, (d) *M. splendida*, (e) *M. strigillosa*, (f) *M. weddelliana*, (g) *M. texana*, and (h) *M. manidea*, shown alongside the first author for size comparison. Images: (a–c) R. F. Barduzzi, (d, g) M. F. Simon, (e–f) L. H. P. Nogueira, (h) L. M. Borges.

**Fig. 2.**
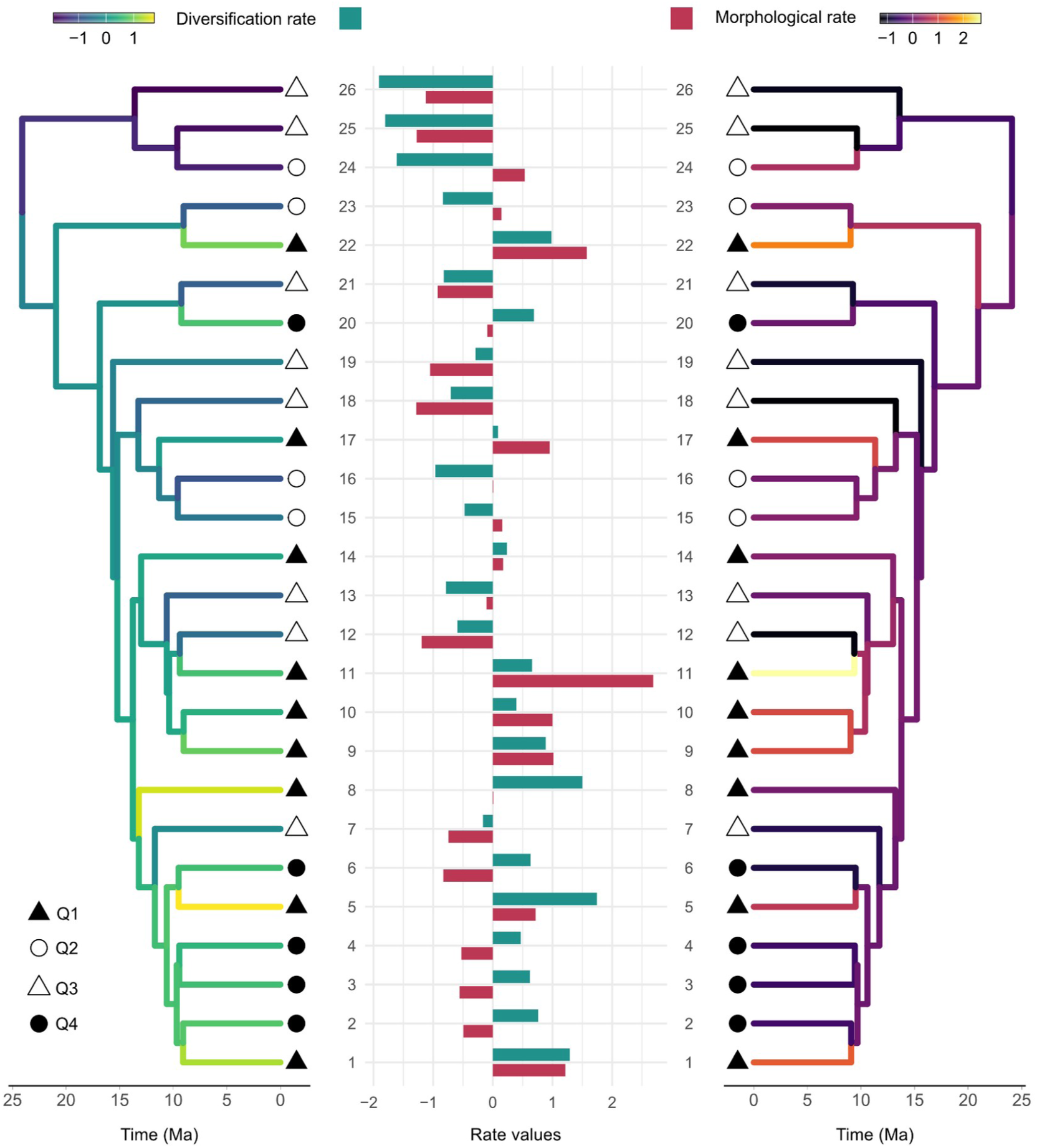
Rates of net diversification and morphological evolution across *Mimosa* clades under the 9 Ma clade delimitation. Net diversification rates plotted here were estimated using the ClaDS model. Both rates were mean-centered and scaled by their standard deviations (and thus have no units). Symbols indicate the radiation-space quadrant to which each clade was assigned based on its diversification and morphological evolution rates. Filled shapes represent radiation-like dynamics and unfilled shapes represent non-radiation-like dynamics. Triangles represent above-average morphological rates, whereas circles represent below-average morphological rates. The bar plots further summarize rates of net diversification and morphological evolution for each clade. Clades with diversification above and below 0 represent the radiation-like (quadrants one and four) and non-radiation-like (quadrants two and three) dynamics, respectively. *Mimosa* taxa belonging to each clade are indicated in Supporting Information Fig. S4.

To understand both the importance of radiations and their drivers at shallow timescales (1–20 Ma; Stephens *et al*., 2025), here we investigate how rates of lineage diversification, morphological evolution, and abiotic niche change are distributed across the phylogeny of *Mimosa*, one of the largest legume genera. Specifically, we ask two questions. First, how much of the biological diversity within *Mimosa* has been generated by different macroevolutionary dynamics? Second, do rates of niche change underlie such dynamics? Given that lineage divergence, all else being equal, provides opportunities for both phenotypic and niche change (Whittaker, 1972; Wills *et al*., 1994), we expected clades with radiation-like dynamics to include the highest proportions of species diversity, morphological disparity, and niche breadth. Furthermore, given that niche lability can facilitate the exploitation of new landscapes or ecological roles, we hypothesized that clades with higher rates of niche change would be more likely to exhibit radiation-like dynamics, whereas those with lower rates would be more likely to exhibit non-radiation-like dynamics (Drummond *et al*., 2012; Stroud & Losos, 2016; Ringelberg *et al*., 2023). Under this hypothesis, variation in rates of abiotic niche change is expected to significantly predict variation in lineage diversification, and perhaps also morphological evolution.

## Materials and Methods

Data processing and analyses were fully conducted using R software version 4.4.0 (R Core Team, 2025), unless stated otherwise.

### Phylogenetic hypothesis and taxonomic units

The most taxonomically comprehensive phylogenies of *Mimosa* currently available were estimated using a single plastid locus, the *trnD–trnT* intergenic spacer (Simon *et al*., 2011; Vasconcelos *et al*., 2020; Ringelberg *et al*., 2023). However, a reanalysis of target-capture data for the tribe Mimoseae, comprising 100 low-copy nuclear loci generated by the large-scale sequencing initiative of Kates *et al*. (2024) across nitrogen-fixing rosid angiosperms, recovered a substantially different backbone for *Mimosa* (Folk *et al*., 2023). To integrate strongly supported nuclear phylogenomic relationships, which include fewer than 25% of the taxa in the genus, with the broader taxonomic sampling available from plastid data (approximately 55% of the taxa in the genus), we adopted a backbone-constrained approach (Supporting Information Fig. S1). This approach was chosen as an alternative to re-estimating backbone relationships from a heterogeneous dataset with incomplete nuclear sampling.

We reanalyzed *Mimosa* plastid data (Simon *et al*., 2011; Vasconcelos *et al*., 2020) in BEAST v.2.7.7 (Bouckaert *et al*., 2014), using a birth–death tree prior and constraining the monophyly of *Mimosa* clades recovered in the Mimoseae phylogenomic tree of Folk et al. (2023). We employed an uncorrelated lognormal relaxed-clock model to accommodate variation in evolutionary rates among branches and used bModelTest to account for uncertainty in nucleotide substitution model selection through Bayesian model averaging (Bouckaert & Drummond, 2017). In the absence of reliable fossil evidence to establish minimum age constraints, we used two secondary calibration points following Simon et al. (2011): the crown node of *Mimosa* and the crown node of *Mimosa* clade A. Both calibration ages were assigned Normal prior distributions, with means of 24.0 and 16.6 Ma, respectively, and standard deviations of 4.0 Ma. Three runs of 50 million generations each were performed, sampling every 10,000 generations. Convergence and stationarity of the MCMC chains were evaluated in Tracer v. 1.7.2 (Rambaut *et al*., 2018). After discarding 10% of the samples from each run as burn-in, the results from the independent runs were combined. Mixing was assessed by examining effective sample sizes (ESS), which generally exceeded 200 for all parameter estimates, as recommended in the BEAST manual. A maximum clade credibility tree was generated from the 13,500 remaining posterior trees.

The current taxonomic delimitation of *Mimosa* includes subspecies and varieties that are currently being either synonymized or elevated to species level (e.g. Santos-Silva *et al*., 2015; Borges *et al*., 2023). To account for differences in taxonomic unit choice, we compared species- and infraspecific-level versions of the same maximum credibility tree in downstream diversification rate analyses.

### Morphological data collection

We compiled data on vegetative morphology for *Mimosa* species from the literature (Barneby, 1991; Du Puy, 1997). We chose to use six leaf traits (leaf-stalk length, pinna rachis length, leaflet length and width, number of pinna pairs, and number of leaflet pairs per pinna; Fig. S2; Table S1) due to the great variation within the group and potential relationships between such variables (e.g. leaf size and shape) and abiotic factors (Yang *et al*., 2015). Such relationships arise from physiological demands such as light capture, thermoregulation, and gas exchange (Givnish, 1979; Reich *et al*., 2003; Wright *et al*., 2004; Reich, 2014; Kraft *et al*., 2015), which can differ enough across a clade to generate conspicuous leaf ecomorphs (e.g. *Viburnum*; Donoghue *et al*., 2022). In addition, we included plant height in our analysis as a proxy for growth form, which is also highly variable and correlated with ecological adaptability within *Mimosa* (Barneby, 1991; Bruneau *et al*., 2024), being recognized as an important trait in the context of plant radiations in general (Givnish *et al*., 2009; Nürk *et al*., 2019). These traits were also widely documented in the literature we searched, resulting in broad phylogenetic sampling (Fig. S3). All trait values were obtained by calculating the midpoint between the lower and higher values reported in the literature.

### Taxa occurrence and abiotic data

We obtained global occurrence data for *Mimosa* from herbarium specimens using GBIF (GBIF, 2020), iDigBio (iDigBio, 2024), and speciesLink (CRIA, 2020). To clean the occurrences, we first removed records lacking species-level identification, not based on preserved herbarium specimens, or containing invalid coordinates. Problematic records were flagged for removal using standardized tests in the ‘CoordinateCleaner’ package (version 3.0.1; Zizka et al., 2019). The package’s recommended settings apply the following criteria to identify problematic records: (1) within a 10,000 m radius of a country’s capital coordinates; (2) within a 1,000 m radius of country centroids; (3) with equal absolute longitude and latitude; (4) within a 100 m radius of biodiversity institutions; (5) located in oceans; and (6) with plain zero coordinates or within a 0.5 m radius of the point 0/0. Finally, all automatically cleaned occurrence records were also curated by specialists, filtering only records that match the native range of taxa.

We updated taxonomic names and corrected typographical errors in taxonomic names using the ‘TNRS’ package (version 0.3.6; Boyle et al., 2013) and the WCVP checklist (Govaerts et al., 2021) as sources. Any name with an overall score equal to or below 0.95 was manually verified again using the GBIF backbone taxonomy database (GBIF, 2024). Afterward, we removed duplicate records and geographical outliers using the respective tests in ‘CoordinateCleaner’ with standard parameters.

Elevation data were obtained from WorldClim 2.1, whereas climatic data were obtained from CHELSA 2.1 (19 BIOCLIM variables, and two BIOCLIM+ variables: the net primary productivity layer, and the mean monthly total cloud cover layer). Edaphic data were obtained from SoilGrids 2.0, averaged across 5-, 15-, and 30-cm sampling depths. We obtained niche predictor values matching the abiotic data to *Mimosa* coordinates. For data analysis, we calculated the median of all values for a given taxon. To reduce multicollinearity, we retained only one predictor from each category (e.g. multiple variables related to air temperature) when variables were highly correlated (Pearson’s |*r*| ≥ 0.85). In such cases, we selected the variable that best represented broad climatic or edaphic conditions and was more directly interpretable in ecological terms, while excluding correlated derivatives or composite variables. The final abiotic dataset comprised elevation, 11 climatic variables, and eight edaphic variables.

### Clade delimitation

All analyses described below required delimitation of clades within *Mimosa*. In order to retrieve clades within a common time frame, we sliced the phylogenetic tree at particular time points using the *treeSlice* function from the ‘phytools’ package (version 2.4-4; Revell, 2024). Since the process of delimiting clades can be somewhat arbitrary (Poe *et al*., 2021), we tried to obtain clades that reflected the differences in morphology and distribution of *Mimosa* using information from both taxonomy and previous delimitations of relevant clades in the context of the diversity and biogeography of the genus (Fig. S4; Barneby, 1991; Simon *et al*., 2011; Vasconcelos *et al*., 2020).

An interval between 11 Ma and 7 Ma matches both the main diversification period of *Mimosa* and the clades previously delimited by Simon *et al*. (2011). Thus, in order to account for the arbitrariness of choosing just one time slice and to explore variation over time in our analysis, we used clade delimitations at 7, 9, and 11 Ma in our analyses, following previous studies (Morinaga *et al*., 2023; Cássia-Silva *et al*., 2026). These three ages (7, 9, 11 Ma) bracket a reasonable number of clades for correlation analyses (35, 26, and 13, respectively; Morinaga *et al*., 2023).

### Rates of net diversification

We estimated net diversification rates for each clade delimitation (7, 9 and 11 Ma) using the Magallón and Sanderson estimator (MS estimator, also known as Method-of-Moments; Magallón & Sanderson, 2001). To estimate rates for clades across each delimitation using this approach, we calculated taxon numbers from taxonomic references and extracted crown or stem ages for each clade. Although the MS estimator assumes a constant relative extinction fraction over time, we can partially account for uncertainty in this fraction by following the standard practice of defining three relative extinction fractions (e.g. 0.0, 0.5, and 0.9; Moen *et al*., 2021).

We also compared these rates (based on taxon diversity and ages) with rates estimated by the cladogenetic diversification rate-shift model (ClaDS; Maliet & Morlon, 2022). We first estimated ClaDS speciation rates for all branches using the same phylogenetic tree and 9 Ma clades used to estimate MS diversification rates. To account for incomplete taxon sampling, we calculated a sampling fraction for each clade by dividing the number of taxa represented in the phylogeny within that clade by the corresponding taxon number used in the MS analyses. Then, we summarized ClaDS output for each clade under the three alternative age delimitations (7, 9, and 11 Ma) by calculating the weighted mean of branch-specific speciation rates, with weights proportional to branch lengths. Finally, we obtained the mean net diversification rate for each clade using the formula: net diversification rate = (1 − ε) * λ_imean_ (Yu & Wiens, 2024).

To evaluate differences when specifying the clade diversity or sampling percentage required in these analyses, we evaluated results considering the number of species when using a species-level phylogeny, or number of infraspecific taxa when using an infraspecific-level phylogeny. All comparisons between diversification-rate estimates and operational taxonomic unit approaches were done using phylogenetic generalized least-squares (pGLS) correlations. We calculated pGLS correlations following Rohlf (2006) and using a custom R script from Moen *et al*. (2013). The estimates of net diversification did not vary significantly between approaches that used either species or infraspecific taxa as operational taxonomic units (Fig. S5). For this reason, we present the results under the species framework from this point onward.

### Rates of morphological evolution and niche change

We estimated multivariate rates of vegetative morphological evolution and rates of niche change (separating altitudinal, climatic, and edaphic niche components) for each clade using the method of Adams (2014), implemented in the ‘geomorph’ package (version 4.0.10; Baken *et al*., 2021). This method calculates a single rate of multivariate, Brownian-motion evolution for multiple characters. In the case of the abiotic niche, we refer to rates of change rather than evolution, because we recognize that factors other than evolution can generate differences in individual niche shifts (Jezkova & Wiens, 2016; Moen & Wiens, 2017). Both morphological and niche predictor data were ln-transformed because phenotypic evolution over such long timescales often occurs multiplicatively, meaning that changes are proportional to trait values rather than absolute (additive) differences (i.e., a doubling of trait value represents an equivalent proportional change regardless of the initial value; O’Meara *et al*., 2006). Moreover, since both types of data also exhibit substantial differences in variance, we mean-centered and scaled the data by their standard deviations prior to the analyses (Adams & Collyer, 2019). We included meristic morphological traits (number of pinna pairs and number of leaflets per pinna) in these transformations because of their approximately log-normal distributions, treating them as pseudo-continuous variables in models of trait evolution.

### Relationships between macroevolutionary rates

Pairwise correlations among rates of diversification, morphological evolution, and niche change were quantified using phylogenetic generalized least-squares (pGLS) correlations. These analyses were exploratory and were used to characterize the magnitude and direction of correlations rather than to evaluate specific biological predictions. We then used pGLS regression with maximum-likelihood estimation of Pagel’s λ, as implemented in the *phylolm* package (version 2.6.5; Tung Ho & Ané, 2014), to explicitly test the a priori hypothesis that rates of niche change predict rates of diversification and morphological evolution. Rates were mean-centered, scaled, and either ln-transformed or left untransformed prior to analysis, depending on their observed distributions and the results of Shapiro–Wilk tests for normality (*n* = 13–35).

### Morphological disparity and niche breadth

We ordinated morphological and niche predictor data and, using independent axes, estimated the breadth of the multidimensional spaces for clades and quadrants using hypervolumes. The ordination was done using phylogenetic PCA as implemented in the ‘phytools’ package. The first five independent axes were used to estimate the hypervolumes using the ‘hypervolume’ package (version 3.1.6; Blonder *et al*., 2018), function *hypervolume_svm*. We chose to calculate hypervolumes on five pPCA axes instead of original data to summarize variation in a reduced number of dimensions (originally seven and 20 variables for morphology and niche, respectively). This approach also allowed us to estimate hypervolumes for clades with as few as five taxa, which was particularly relevant given that five of the 26 clades at 9 Ma had only five taxa due to low diversity or limited data.

### Radiation space

Following the approach of previous studies, we define a radiation space using quadrant analysis within a rate-defined space (Sanderson, 1998; Lieberman, 2012; Moen *et al*., 2021; Morinaga *et al*., 2023). Clades with lineage diversification rates above the mean were classified as radiation-like (two right-hand quadrants), whereas those below the mean were classified as non-radiation-like (two left-hand quadrants). Radiation-like clades in the first (upper-right) quadrant also exhibit rates of morphological evolution above the mean, while non-radiation-like clades in the third (lower-left) quadrant exhibit rates of morphological evolution below the mean. We chose to omit the term ‘adaptive’ when referring to radiation types because, although our trait selection prioritized plant traits with presumed ecological importance, no explicit form–function analysis was conducted (Givnish, 2015; Moen *et al*., 2021). The occurrence of a clade in a quadrant is here interpreted as a hypothesis about its overall macroevolutionary dynamic (Morinaga *et al*., 2023).

## Results

### Macroevolutionary rates

The estimates of net diversification using the ClaDS model and the MS estimator showed somewhat different results. Under the 9 Ma clade delimitation, correlations between ClaDS and MS ranged from 0.62 to 0.81 under different extinction values when using crown ages, and from 0.69 to 0.70 when using stem ages. The rates for clades varied approximately 2.5-fold, from 0.176–0.436 net diversification events per million years in the ClaDS results, approximately 8-fold, from 0.072–0.560 net diversification events per million years in the MS results when using crown ages, and approximately 8-fold, from 0.051–0.398 net diversification events per million years when using stem ages. These values place *Mimosa* diversification rates below those reported for Andean *Lupinus* and Campanulaceae (Hughes & Eastwood, 2006; Lagomarsino *et al*., 2016), but within the range reported for *Viburnum* and Andean *Macrocarpaea* (Spriggs et al., 2015; Vieu et al., 2022). For the sake of simplicity, we mainly show and discuss diversification rates estimated using ClaDS from this point onward, while also considering results from the MS estimator (ε = 0.5) based on either crown ages (MS-crown) or stem ages (MS-stem) when discrepancies arise. Across clades, rates of morphological evolution varied >67-fold, from 0.004 to 0.262 composite trait variance per million years.

### Relationship between rates

The pGLS regression revealed significant positive associations between the rate of diversification and both edaphic and altitudinal niche change under the 7 Ma clade delimitation (Table 1). This pattern was also observed under the 9 Ma delimitation, although only for MS-crown (Table S3). All the rates of niche change (altitudinal, edaphic, and climatic) were associated with morphological evolution under the 7 Ma clade delimitation, but not under the other delimitations (Table 1; Table S4). Under the older, 11 Ma delimitation, no rate of niche change emerged as a significant predictor of either diversification or morphological evolution (Tables S2, S3, and S4). Results also indicate that altitudinal niche change and edaphic niche change yield greater explanatory power (R²) over rates of diversification and morphological evolution, respectively (Table 1). Although rates of climatic niche change appear as a predictor of rates of morphological evolution, their explanatory power (R²) is weaker (Table 1).

**Table 1.** pGLS analyses of ClaDS diversification rates and morphological evolution rates as a function of niche-change rates under the 7 Ma clade delimitation. *P*-values are two-sided, based on Wald tests of the regression coefficients. \**P* < 0.05.

| Model | Slope | Adjusted *R*<sup>2</sup> | *P*-value |
|---|---|---|---|
| Diversification rate ~ Altitudinal rate | 0.390 | 0.170 | 0.008* |
| Diversification rate ~ Climatic rate | 0.151 | 0.006 | 0.281 |
| Diversification rate ~ Edaphic rate | 0.341 | 0.137 | 0.016* |
| Morphological rate ~ Altitudinal rate | 0.452 | 0.180 | 0.006* |
| Morphological rate ~ Climatic rate | 0.413 | 0.145 | 0.014* |
| Morphological rate ~ Edaphic rate | 0.520 | 0.248 | 0.001* |

### The radiation space of *Mimosa*

As indicated by the distribution of clades in the radiation space, we found a moderate correlation between rates of diversification and morphological evolution (*r* = 0.555, *P* = 0.049, *n* = 26; Fig. 3). Nonetheless, clades appear to be approximately evenly distributed in the radiation space, with 14 radiation-like clades and 12 non-radiation-like clades. At least 70% of disparity and diversity were attributed to radiation-like dynamics under the 9 Ma clade delimitation, with first-quadrant radiations accounting for the largest contribution (>65% for both metrics; Fig. 3). The remaining diversity and disparity were attributed to non-radiation-like dynamics (around 10–20% for both metrics; Fig. 3; Fig. S6). Under the younger, 7 Ma clade delimitation, this pattern was largely maintained, although the contribution from non-radiation-like dynamics to diversity and disparity had generally increased (Fig. S6). Under the older, 11 Ma clade delimitation, the contribution of radiation-like dynamics generally increased, with at least 80% of disparity and diversity being attributed to these dynamics, particularly first-quadrant radiations (Fig. S6). While results from MS-crown agree with the values and patterns presented above, MS-stem results do not show a clear pattern of radiation-like dynamics increasing their contribution to diversity and disparity across time (Fig. S6).

**Fig. 3.**
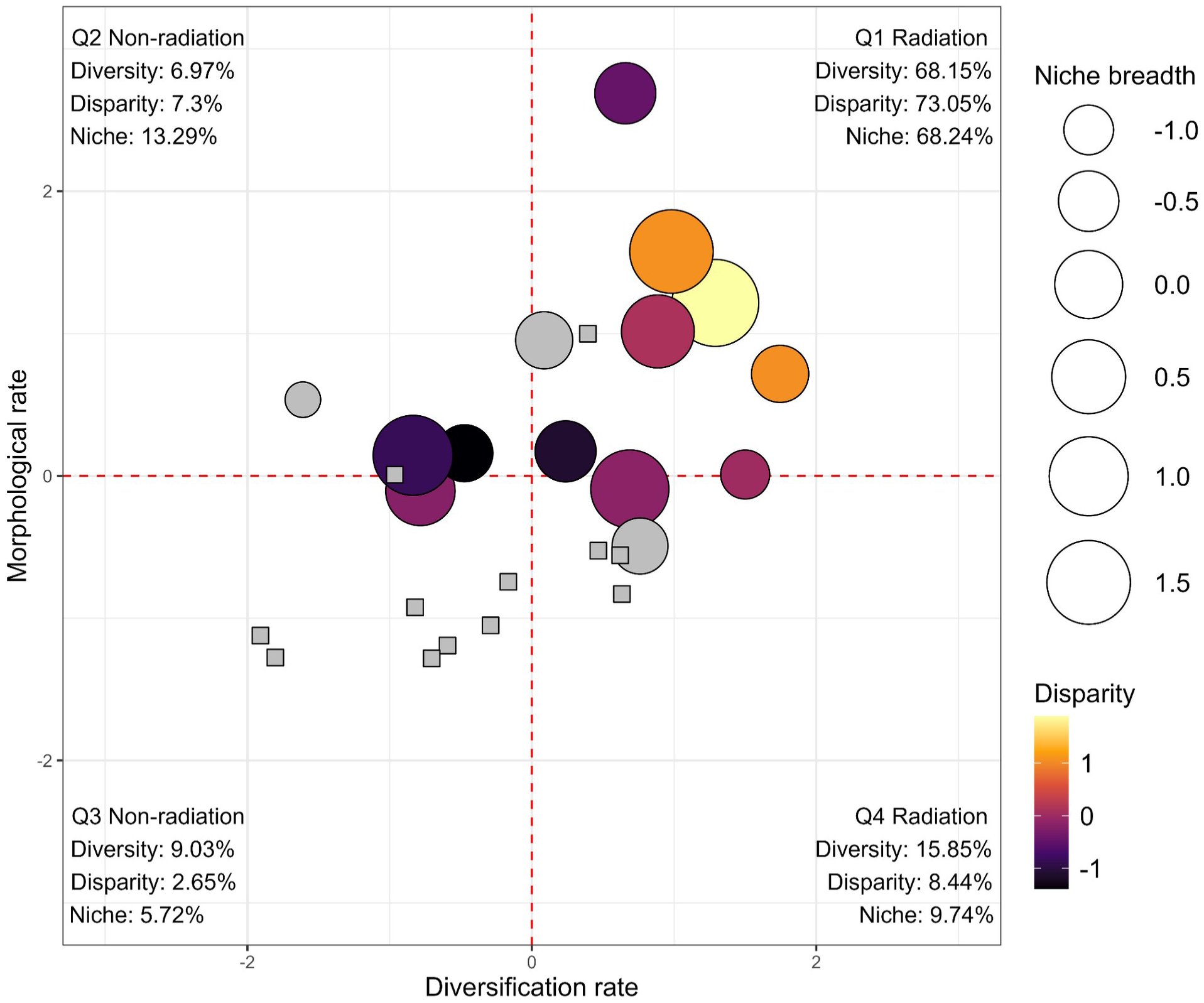
The radiation space of *Mimosa* under the 9 Ma clade delimitation. Dots show the estimated rates of diversification and morphological evolution for each *Mimosa* clade (*n* = 26). Both rates were mean-centered and scaled by their standard deviations. Percentages indicate the proportion of the diversity, disparity, and niche breadth of *Mimosa* associated with each quadrant (macroevolutionary dynamic). Disparity and niche breadth were both assessed using a five-dimensional hypervolume. Percentages of disparity and niche breadth across quadrants may sum to >100% or <100% because of overlaps or gaps, respectively, between quadrants in morphospace and niche space. Dot size and color indicate niche breadth and the magnitude of morphological disparity, respectively, for each clade. Gray dots indicate clades for which disparity could not be estimated because of insufficient sampling (*n* < 6 species), whereas gray squares indicate clades for which neither metric could be estimated for the same reason.

Similar to disparity and diversity, at least 60% of the niche breadth of *Mimosa* (including all niche components) was generally attributed to radiation-like dynamics under the 9 Ma clade delimitation (Fig. 3). However, results from MS-crown attribute 45% of this metric to non-radiation dynamics under the same clade delimitation (Fig. S6). Under the younger, 7 Ma clade delimitation, around 5–25% of the niche breadth was attributed to non-radiation-like dynamics, similar to what is observed for diversity and disparity (Fig. S6). Under the older, 11 Ma clade delimitation, at least 80% of niche breadth was commonly attributed to radiation-like dynamics. However, for the MS-stem results, non-radiation-like dynamics seem to account for most of the niche breadth (Fig. S6).

## Discussion

Here, we evaluated whether radiation-like dynamics underlie most biological diversity in one of the largest genera of legumes, and the potential role of abiotic niche change in promoting such dynamics. We found that at least 70% of disparity and diversity are consistently attributed to radiation-like dynamics in clades delimited using a 9 Ma age threshold, whereas 45% of niche breadth is sometimes attributed to non-radiation-like dynamics at the same timescale (Fig. 3; Fig. S6). Moreover, under the 7 Ma clade delimitation, variation in diversification rates was positively associated with altitudinal and edaphic niche change across clade delimitations, whereas morphological evolution was positively associated with all rates of niche change (Table 1). Finally, for most methodological scenarios, the contribution of radiation-like dynamics to biodiversity appears to increase under the 11 Ma clade delimitation (Fig. S6).

### Macroevolutionary dynamics underlying *Mimosa* diversity

Our results indicate that radiation-like dynamics, particularly those accompanied by above-average rates of morphological evolution, account for a substantial proportion of species diversity and disparity in *Mimosa* (Fig. 3). Although not explicitly quantified previously, available evidence suggests similar inferences in other plant groups with relatively recent diversification histories (∼30–20 Ma). Within Leguminosae, a previous study on the genus *Lupinus*, for example, indicated that most of its estimated extant species diversity is concentrated in clades classified as recent, rapid radiations (Drummond *et al*., 2012). Similarly, rapid island radiations have been associated with the majority of growth-form disparity across distantly related plant genera (Nürk *et al*., 2019). However, accelerated diversification does not always need to be the main dynamic underlying such biodiversity metrics. This may be particularly true for plant lineages from regions with relatively stable climates, where radiation events are hypothesized to play a minor role compared to the gradual accumulation of lineages (Nge & Skeels, 2025). In the old, climatically buffered Southwest Australian Biodiversity Hotspot, for example, the *Cassinia–Ozothamnus* (Asteraceae) clade diversified within the last 15 Ma and most of its species diversity does not seem to be tied to the single shift in diversification rate inferred for the clade (Nge & Skeels, 2025).

As revealed by these contrasting examples, associations between biodiversity metrics (e.g. diversity and disparity) and their related rates (e.g. diversification and morphological evolution), although expected, are not guaranteed. Both simulation and empirical research have revealed that the correlation between diversity and diversification rates can be weak or negative, especially at shallow timescales such as the one evaluated here (Kozak & Wiens, 2016; Dimitrov *et al*., 2023; Stephens *et al*., 2025). This pattern can emerge, for example, in scenarios where new ecological opportunities were accessed relatively recently in evolutionary history (Stephens *et al*., 2025) yet insufficient time has passed for diversity to relate to diversification rates (Stephens & Wiens, 2003). Similarly, the relationship between disparity and rates of morphological evolution can vary across lineages, with some generating different morphologies through the rapid recombination of existing trait states (Felice *et al*., 2018), resulting in an exhaustive ‘running in circles’ pattern in morphospaces (Santos *et al*., 2019; Barduzzi *et al*., 2026). These examples provide a useful framework for interpreting the patterns observed in *Mimosa*, in which a relatively smaller proportion of diversity and disparity is underlain by above-average rates of diversification and morphological evolution, respectively, compared with previous studies (Morinaga *et al*., 2023; Wiens & Moen, 2025).

Overall, although diversity and disparity appear closely associated with radiation-like dynamics, *Mimosa* clades show a relatively balanced distribution across the radiation space (i.e., an approximately even split between radiation-like and non-radiation-like clades; Fig. 3), indicating a plural history of macroevolutionary dynamics across the phylogeny. This matches expectations based on the diversity and wide geographic distribution of the genus (Barneby, 1991; Simon *et al*., 2011; Borges *et al*., 2024). Such a plural evolutionary history, which may not be common in other highly diverse plant genera with more uniform morphology and niche biology (Folk *et al*., 2024), makes *Mimosa* an important model for future work seeking to identify factors behind contrasting dynamics of diversification, morphological evolution, and niche change. Furthermore, elevated disparity and/or niche breadth make some clades within or close to the first quadrant (e.g. clades 1, 5, and 22; Fig. S7) interesting candidates for evaluating adaptive morphological evolution. This aspect could be evaluated, for example, through comparative, functional, or population-level analyses, including an assessment of the repeated evolution of ecologically analogous taxa (Givnish, 2015; Moen *et al*., 2021; Donoghue *et al*., 2022).

Our evaluation of the importance of radiation-like dynamics to niche evolution, which treated this aspect of biodiversity independently rather than as part of ecomorphological variation, as commonly done for animals (e.g. Huie *et al*., 2021; Morinaga *et al*., 2023), shows that radiation-like dynamics do not always clearly account for most of *Mimosa*’s niche breadth (Fig. S6). This suggests that radiation-like dynamics do not necessarily unfold across a broad range of environments but can be instead restricted to particular abiotic niches. Links between abiotic niche conservatism and high diversification rates have been explored previously (Wiens *et al*., 2010; Hua & Wiens, 2013), including in other plant groups (Folk *et al*., 2019). This relationship can arise, for example, when climatic niche conservatism promotes the isolation of populations that may then undergo allopatric speciation (Kozak *et al*., 2006; García-Navas & Westerman, 2018; Qiao *et al*., 2024). Thus, in *Mimosa*, a high contribution of slower diversification dynamics to abiotic niche breadth in some of our results may indicate that niche exploration can unfold relatively independently of lineage diversification. Rather than being a general consequence of expanding into new abiotic niches, radiation-like dynamics may instead be associated with niche lability under particular environmental conditions.

### In the right places: the role of niche lability in *Mimosa* diversification history

Our hypothesis that niche lability generates opportunities that can promote radiation-like dynamics was strongly supported at least in the most recent timescale (Table 1). Although rates of climatic niche change did not predict diversification, we recovered a positive relationship between rates of lineage diversification and rates of both altitudinal and edaphic niche change under the 7 Ma clade delimitation (Table 1). Moreover, rates of altitudinal, climatic, and edaphic niche change were positively associated with rates of morphological evolution under the same clade delimitation (Table 1). This indicates that more labile lineages, which navigate niche space more rapidly (though not necessarily across a broader range of niches), also diversify—and occasionally evolve across trait space—more rapidly.

The link between both rates of diversification and morphological evolution with altitudinal niche change indicates that parapatric and allopatric dynamics may play a role in radiation-like dynamics across the broad range of montane habitats in which *Mimosa* is particularly diverse (Simon *et al*., 2011; Cadena *et al*., 2011; Pyron *et al*., 2015). Specifically, the rapid partitioning of altitudinal niches observed in some *Mimosa* clades could be explained by populations occasionally shifting away from their ancestral altitudinal niche and subsequently tracking their newly acquired niche, thereby reducing gene flow between populations (Pyron *et al*., 2015). Such a scenario may be further reinforced by the usually short-distance dispersal mechanisms of montane plant lineages (Morgan & Venn, 2017; Ribeiro *et al*., 2023; Di Musciano *et al*., 2023). For example, in *Navia* Schult. & Schult.f. (Bromeliaceae), which is particularly diverse in the Guiana Highlands, high species diversity has been associated with large, naked seeds that are dispersed over short distances (Givnish, 1997, 2015). Similarly, dispersal limitation may be an important factor underlying the radiation of Andean *Lupinus* (Contreras-Ortiz *et al*., 2018). Despite a few exceptions, the majority of *Mimosa* species lack specialized mechanisms for long-distance dispersal (Barneby, 1991), with seeds tending to drop near parental plants (personal observation in the field). Thus, the combination of limited dispersal and heterogeneous landscape opportunities might have triggered independent *Mimosa* radiation-like events across altitudinal gradients.

In montane floras, edaphic niche lability is also expected to foster diversification, particularly given the environmental heterogeneity within mountains and between mountains and lowlands (Körner, 2002; Hughes & Eastwood, 2006; Drummond *et al*., 2012; Hughes & Atchison, 2015; Cortés & Wheeler, 2018). As physical and chemical soil conditions largely determine nutrient and water availability, pronounced edaphic heterogeneity can alter fundamental processes underlying the evolution of plant lineages, including lineage diversification and ecological divergence (Rajakaruna, 2018; Hulshof & Spasojevic, 2020). Indeed, edaphic heterogeneity may foster ecological divergence—often reinforced by restricted gene flow through shifts in flowering time (Rajakaruna, 2018)—even between geographically adjacent populations (e.g. Cabrini *et al*., 2025). Moreover, divergence in vegetative traits such as plant stature and leaf dimensions is a relatively common response to physiological stress related to edaphic transitions (Alcantara *et al*., 2015; Hulshof & Spasojevic, 2020; Samojedny *et al*., 2022). This pattern may explain the further link between edaphic niche change and rates of morphological evolution uncovered here. The topographical heterogeneity along the altitudinal gradient of mountains amplifies the effects of soil variation in reducing gene flow between populations, thereby increasing diversification rates over evolutionary time (Pyron *et al*., 2015; Hulshof & Spasojevic, 2020; Tietje *et al*., 2022).

Taken together, our results are consistent with a scenario in which both geographic and ecological divergence play key roles in driving radiation-like dynamics, albeit through contrasting mechanisms. Shifts in altitudinal niche may have fostered topographic isolation among populations distributed across distinct altitudinal belts, representing a non-adaptive geographic component. Conversely, edaphic niche changes may have fostered ecological divergence among populations, representing an adaptive dimension of the radiation-like dynamics recovered here for *Mimosa* (Gittenberger, 1991; Givnish, 2015; Donoghue *et al*., 2022). However, not all abiotic niche dimensions are associated with radiation-like dynamics, given that rates of climatic niche change are only linked to rates of morphological evolution, even though they are partially correlated with the rates of altitudinal niche change (*r* = 0.689, *P* < 0.001, n = 26). The link between climatic niche change and morphological rates could have arisen through processes of adaptation in the vegetative morphology, such as adaptation to different precipitation regimes (Wang *et al*., 2022), that unfolded relatively independently of lineage diversification (Folk *et al*., 2019; Li *et al*., 2024). Overall, although both lability and conservatism in climatic niche may contribute to isolation along altitudinal gradients (Qiao *et al*., 2024; Dellinger *et al*., 2024), changes in niche dimensions more directly associated with the physical environment (i.e., elevation and soil), which persist through geological time (Corlett & Tomlinson, 2020), are more strongly correlated with variation in both rates of lineage diversification and morphological evolution in *Mimosa*.

Finally, although our results indicate that niche change at a coarse spatial scale plays a prominent role in *Mimosa* diversification, abiotic niche variation at finer spatial scales than those quantified here may also be important. For instance, the Campos Rupestres of central Brazil, one of the diversity hotspots of the genus (Barneby, 1991), includes a wide range of microhabitats not captured in our analyses (e.g. local variation in soil texture and coarseness; Alves *et al*., 2014; Barros-Souza *et al*., 2026). Some of the clades occurring in the Campos Rupestres (e.g. clades 7 and 9, corresponding to clades O and R of Simon *et al*. 2011, in Fig. S4) feature radiation-like dynamics but show very low rates of niche change, even along the altitudinal axis (Fig. S8). Although dynamics of isolation-driven divergence due to altitudinal niche conservatism could explain such a contrasting scenario in the Campos Rupestres (e.g. Verboom *et al*., 2015; Vasconcelos *et al*., 2020), local variation in edaphic conditions not captured here likely plays an important role in lineage divergence (e.g. Cabrini *et al*., 2025). Future research assessing the role of finer-scale environmental heterogeneity on macroevolutionary rates may elucidate the role of microhabitats in driving radiation-like dynamics. Moreover, explicitly accounting for the geographic distribution of lineages (e.g. the effects of range breadth, expansion, and overlap on diversification rates; Nery *et al*., 2026) could help to elucidate the hypotheses we raised here concerning the role of biogeography in the diversification history of *Mimosa*. Nonetheless, we argue that topographically heterogeneous regions, including major centers of *Mimosa* diversity (e.g. the Mexican, Malagasy, Andean, and Central Brazilian highlands; Barneby, 1991; Simon *et al*., 2011; Barros-Souza & Borges, 2023), acted as key landscapes (*sensu* Givnish, 1997). By providing physiographically heterogeneous settings, these regions promoted independent radiation-like dynamics within the genus (Bouchenak-Khelladi *et al*., 2015; Givnish, 2015; Hughes *et al*., 2015).

### Do radiations underlie most of biodiversity at shallower timescales?

Events of rapid lineage diversification, here regarded as ‘radiations,’ are thought to underlie much of life’s diversity, especially at higher taxonomic scales such as families and orders (Simpson, 1953; Simões *et al*., 2016; Wiens & Moen, 2025). For frog families, both diversity and disparity seem to match this expectation (Morinaga *et al*., 2023). Similar evidence indicates that the species diversity of plant phyla is also mostly underlain by radiations (Wiens & Moen, 2025). However, as previously seen for some groups of animals (Morinaga *et al*., 2023; Stephens *et al*., 2025), here we show that despite accounting for a substantial proportion of lineage diversity and morphological disparity, the contribution of rapidly diverging lineages to the generation of biodiversity tends to weaken at shallower timescales (Fig. S6).

Although current evidence is insufficient to determine if this temporal pattern of radiations is common, its occurrence in widely different groups indicates it may be widespread across the tree of life. We propose two explanations for this pattern. First, it may be harder to discern the contribution of rapidly diversifying lineages to biodiversity closer to the present because these lineages may not yet have had sufficient time to accumulate a large number of descendants, despite their high diversification rates (Stephens and Wiens, 2003). Consequently, young lineages undergoing rapid diversification may not yet exhibit the relatively high species diversity typically associated with radiations, making them more difficult to identify as such. This possibility is consistent with simulations showing that the accuracy of diversification-rate estimates increases strongly with clade age, such that rates estimated for young clades may provide less accurate measures of their diversification dynamics (Kozak & Wiens, 2016). Second, time is important for sorting the products of different macroevolutionary dynamics, which would be more diverse near the present. As time accumulates, so do the effects of increased diversification rates, leading to the overrepresentation of such lineages—even if due to the increased extinction likelihood of smaller clades (Jablonski, 2017)—in the composition of extant biodiversity (Pontarp & Wiens, 2017; Stephens *et al*., 2025). As these factors are not mutually exclusive, the increasing contribution of radiations at deeper timescales may well result from the combined effects of methodological accuracy to detect patterns of diversification as well as the accumulation of biodiversity by lineages that are able to rapidly generate new lineages.

Despite their time-dependent but significant role in the generation of diversity and disparity, radiations do not clearly underlie most of the abiotic niche breadth occupied by *Mimosa* (Fig. S6). This result suggests that some radiations may occur under specific environmental sets (as discussed above), such as radiations dominated by allopatric speciation or by differentiation along microhabitat rather than macrohabitat dimensions (Rundell & Price, 2009; Wiens *et al*., 2010). In these cases, niche conservatism at larger environmental scales can be connected to increased diversification (Pyron *et al*., 2015; Qiao *et al*., 2024). For example, in Hawaiian arthropods, the niche in a macro-scale is conserved in adaptive radiations relative to microhabitat preferences (Hiller *et al*., 2019). On the other hand, for some groups (e.g. *Viburnum* L., Adoxaceae; Donoghue *et al*., 2022), changes in both macro- and microclimatic niches are linked to repeated radiations across space. Thus, understanding the potential importance of the detailed, fine-grained abiotic niche (Hiller *et al*., 2019; Donoghue *et al*., 2022), a dimension not explored here, is also important for disentangling the role of niche change across different macroevolutionary dynamics.

Overall, although the idea that radiations underlie most biodiversity has so far been supported (Morinaga *et al*., 2023; Wiens & Moen, 2025), deviations from this expectation across groups, scales, and metrics—such as the one we highlighted for niche breadth—are likely to improve our understanding of diversification dynamics. For example, it is possible that not only niche breadth but also disparity may not be clearly underlain by radiations in other groups, as suggested by the decoupling of taxonomic diversity and disparity, even under such macroevolutionary dynamics (Wills *et al*., 1994; Ruta *et al*., 2013; Oyston *et al*., 2016). Finally, we echo the call for additional quantitative tests of the role of different macroevolutionary dynamics in the generation of biodiversity. Frameworks like the one used here (Morinaga *et al*., 2023; Wiens & Moen, 2025), which merges categorical classifications of macroevolutionary dynamics and continuous variation in rates of evolution, are promising candidates for investigating how a continuum of evolutionary patterns and processes (Olson & Arroyo-Santos, 2009) shapes biological diversification.

### Conclusions

As a whole, our work highlights that both relatively rapid and slow lineage diversification can underlie the multifaceted nature of biodiversity. We show that the generation of a substantial proportion of species diversity and morphological disparity was driven by radiation-like dynamics in *Mimosa*, a highly diverse plant genus. However, in contrast to previous findings, slower diversification dynamics not only underlie part of the patterns of diversity and disparity but could also account for a substantial portion of the genus’s niche breadth. Moreover, a coupling between rates of niche change and both diversification and morphological evolution suggests that increased biological diversification is associated with niche lability and spatial heterogeneity, reinforcing the importance of these axes for plant macroevolution. More broadly, the application of an emerging framework to study diversification at a relatively shallower time frame is a further step toward quantifying the contribution of different macroevolutionary dynamics to the generation of biological diversity across organisms and evolutionary time. Finally, we emphasize that biodiversity is multifaceted and argue that this complexity must be embraced if we are to fully understand how variation in macroevolutionary tempo, beyond a simple tortoise-and-hare scenario, gives rise to the diversity of life.

## Acknowledgements

We thank the members of the Taxonomy and Evolution of Plants lab (Universidade Federal de São Carlos, Brazil) and Moen lab (University of California, Riverside, USA) for discussions during the development of this research. We also thank Monique Maianne for her support in morphological analyses and for providing valuable comments at different stages of the research. This work was supported by the Coordenação de Aperfeiçoamento de Pessoal de Nível Superior (CAPES; Finance Code 001), the Fundação de Amparo à Pesquisa do Estado de São Paulo (FAPESP; grant numbers 2024/20286-0 and 2023/17978-5 to R.F.B., 2023/16170-4 and 2021/13031-8 to Y.B.-S., and 2022/03046-0 to L.M.B.), and the U.S. National Science Foundation (NSF; grant IOS-2430681 to D.S.M.), which supported R.F.B.’s participation in a workshop on phylogenetic comparative biology.

## Author contributions

RFB, LMB, and DSM conceptualized the study. RFB, YB-S, and LMB collected and curated the data. RFB conducted all data analyses. RFB wrote the first draft; all authors revised and approved the manuscript. LMB and DSM contributed equally to this work.

## Data availability

The data and code supporting the findings of this study will be made publicly available upon publication.

## Supporting Information

Additional Supporting Information may be found online in the Supporting Information section at the end of the article.

**Fig. S1** Topological differences between *Mimosa* phylogenies estimated using different methodological approaches.

**Fig. S2** Leaf traits evaluated.

**Fig. S3** Phylogenetic sampling of vegetative data.

**Fig. S4** *Mimosa* taxonomy and clade delimitations.

**Fig. S5** Correlation among diversification rate estimation methods under the 9 Ma clade delimitation.

**Fig. S6** Comparison of *Mimosa* radiation spaces using different diversification estimators and clade delimitations.

**Fig. S7** The radiation space of *Mimosa* under the 9 Ma clade delimitation.

**Fig. S8** Rates of net diversification and altitudinal niche change across the *Mimosa* phylogeny under the 9 Ma clade delimitation.

**Table S1** Vegetative morphological traits compiled from the literature.

**Table S2** pGLS analyses of ClaDS diversification rates as a function of niche change rates across clade delimitations.

**Table S3** pGLS analyses of Magallón and Sanderson estimator diversification rates as a function of niche change rates across clade delimitations.

