## Supporting information for "Beyond tortoises and hares: Niche lability and a continuum of diversification dynamics underlie the megadiversity of a legume genus"

The following Supporting Information is available for this article:

### Supplementary Figures

**Fig. S1: Topological differences between *Mimosa* phylogenies estimated using different methodological approaches.** The phylogeny on the left was estimated in this study using plastid data constrained by phylogenomic information, while the one on the right was reported by Vasconcelos *et al.* (2020), based exclusively on plastid data.

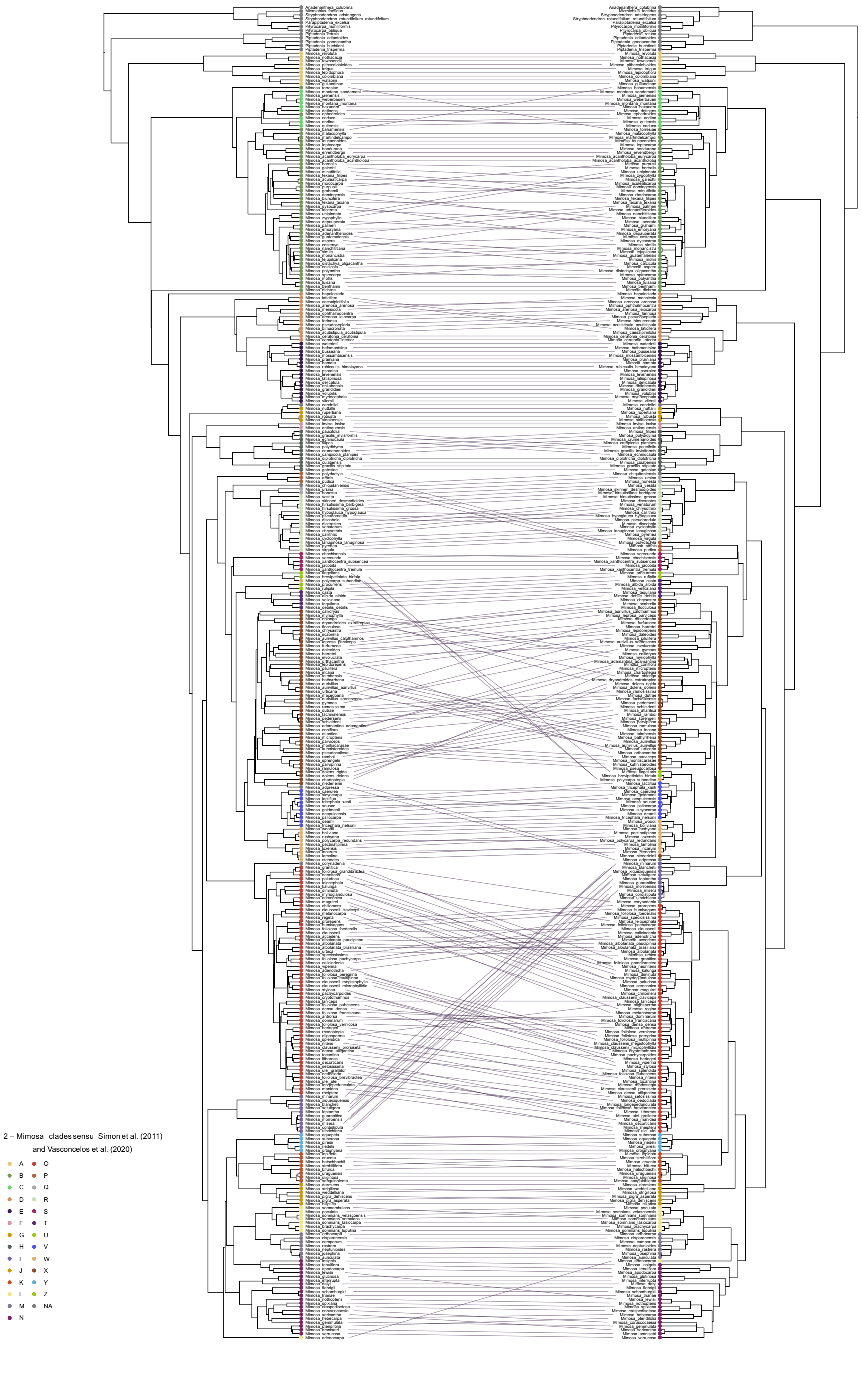

**Fig. S2: Leaf traits evaluated.** The darker dots represent the pulvinated regions. Leaflets and pinnae were counted in pairs. The leaf illustrated is from *Mimosa candollei*. Image: R. F. Barduzzi.

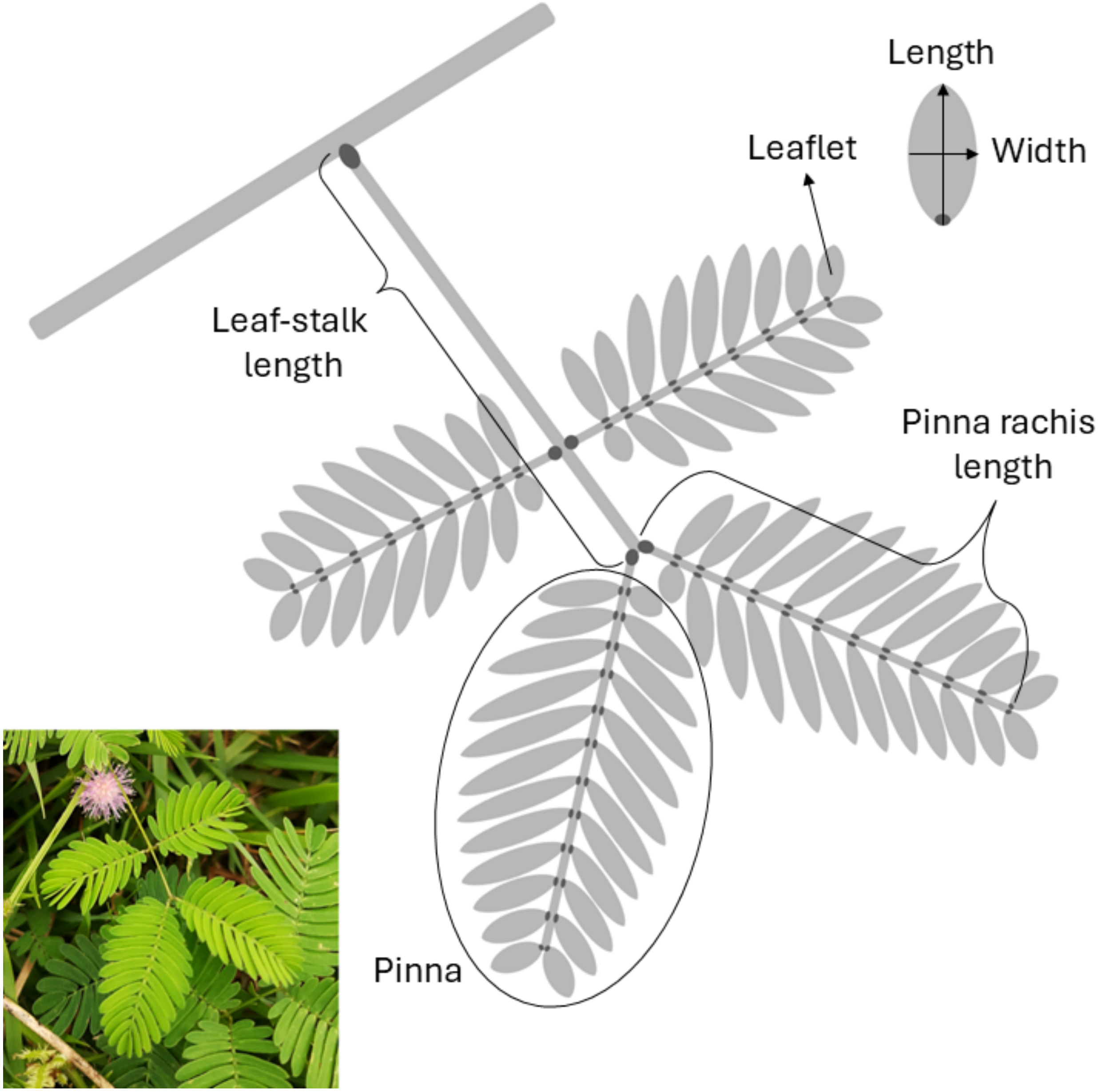

**Fig. S3: Phylogenetic sampling of vegetative data.** Only *Mimosa* taxa with complete descriptions of the selected traits were included in our analyses.

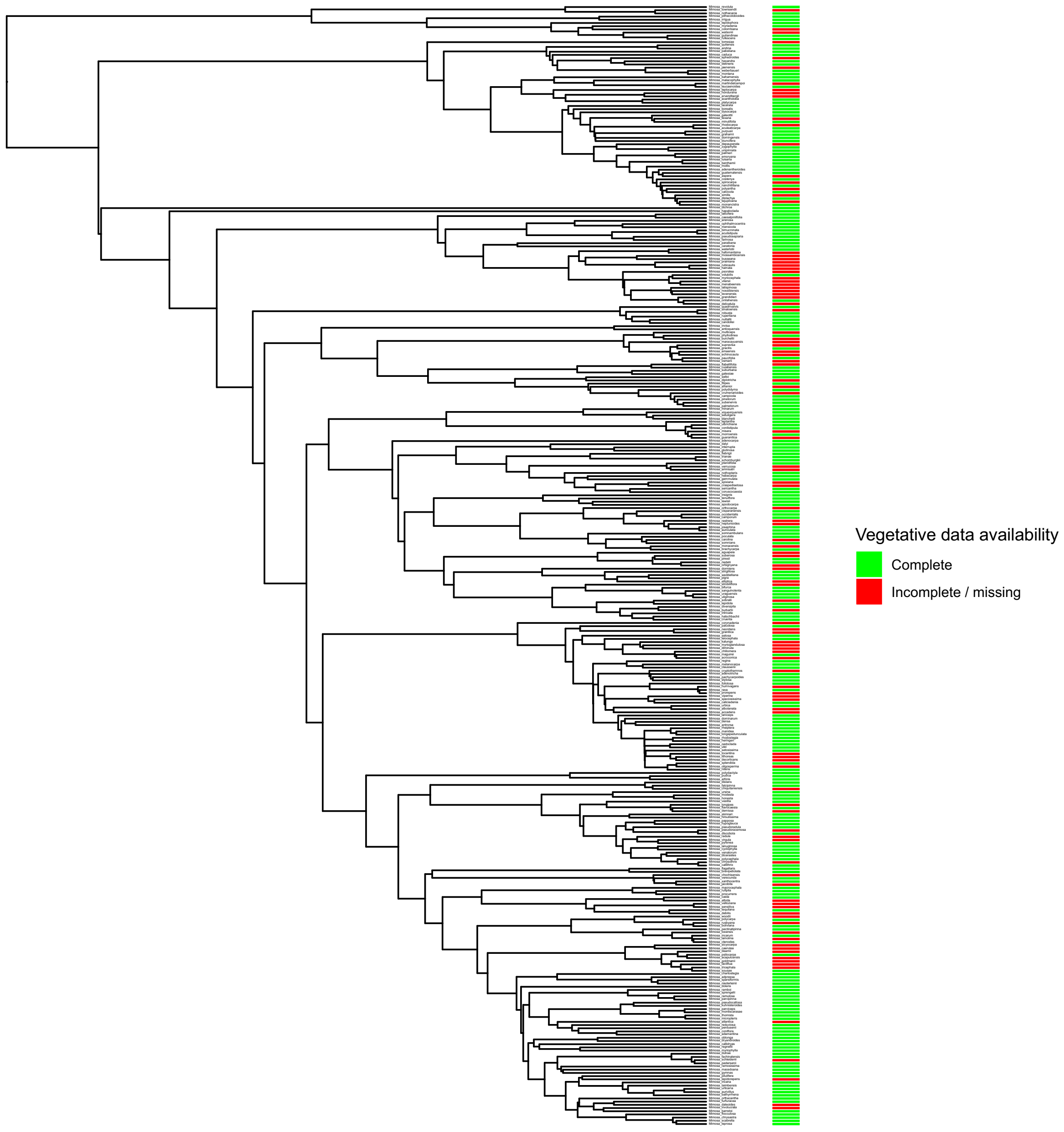

**Fig. S4: *Mimosa* taxonomy and clade delimitations.** The heatmap shows a clade delimitation at 9 million years ago (Ma), the non-monophyletic taxonomic treatment of *Mimosa sensu* Barneby (1991), and previous clade delimitations proposed by Simon *et al.* (2011) and Vasconcelos *et al.* (2020).

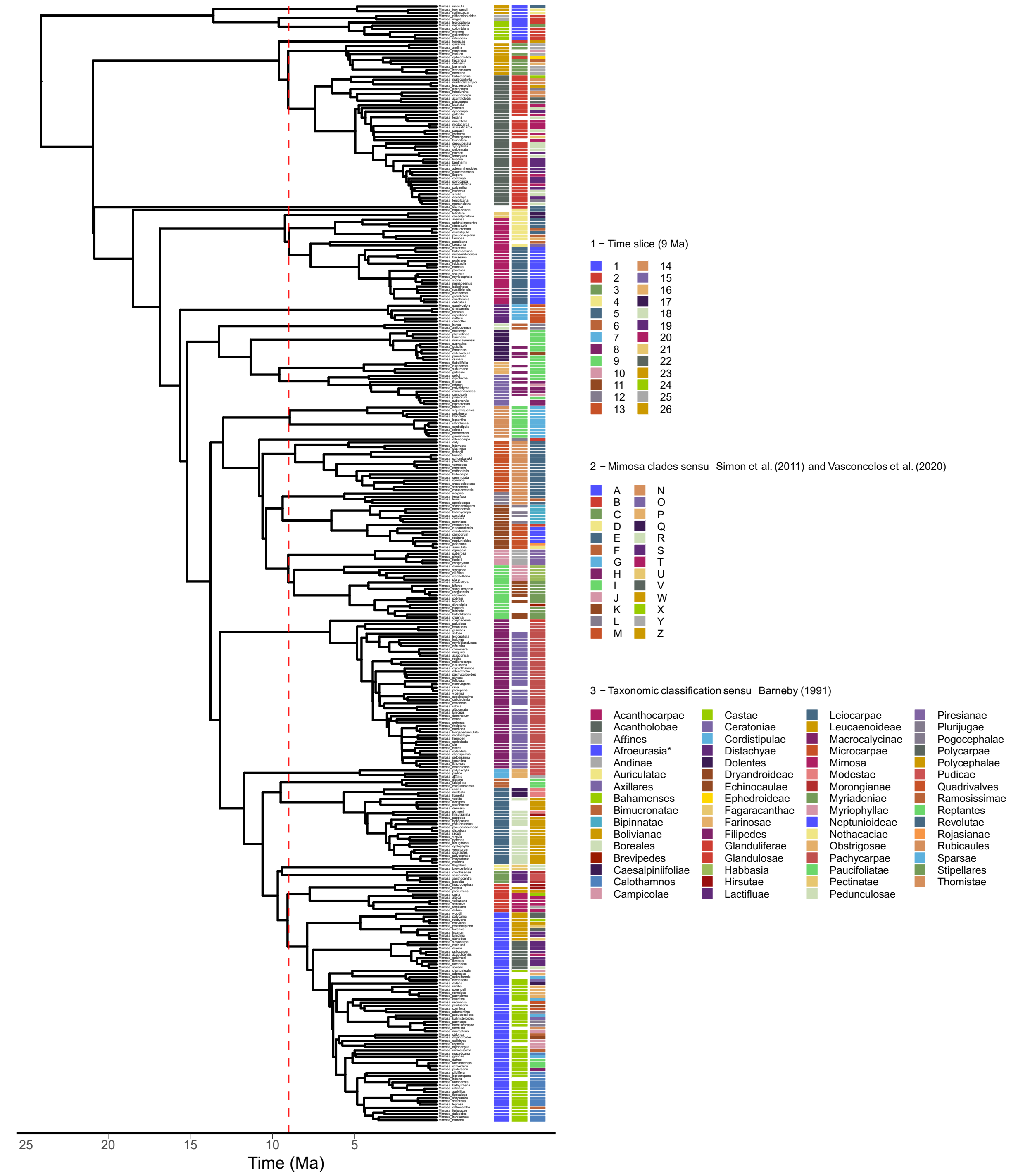

**Fig. S5: Correlation among diversification rate estimation methods under the 9 Ma clade delimitation.** We compared the Magallón and Sanderson estimator (MS) results, using both crown and stem ages and different values of extinction (low, medium, and high), with the ClaDS model results. We also estimated diversification rates under two scenarios: considering only species-level taxa (sp) or including all infraspecific taxa (ifsp).

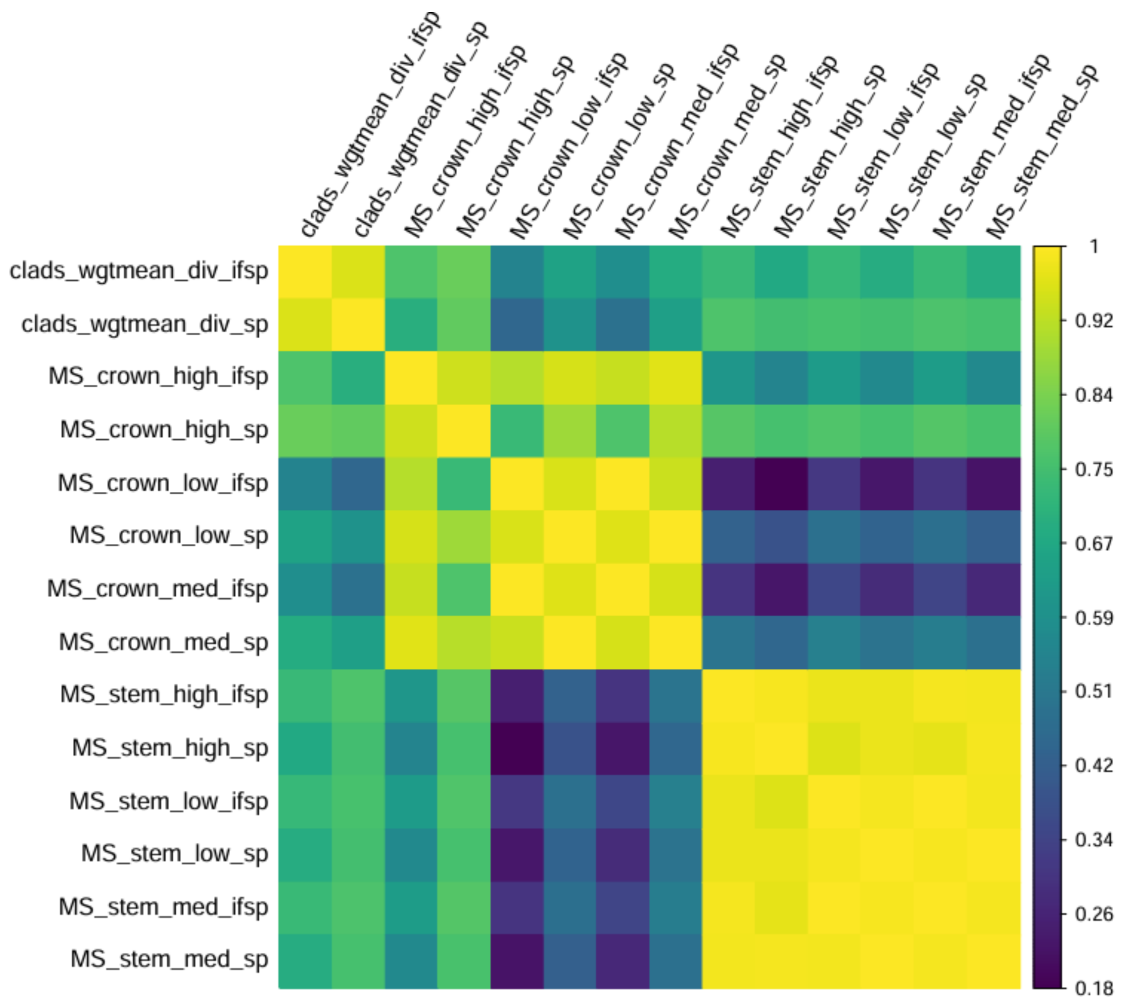

**Fig. S6: Comparison of *Mimosa* radiation spaces using different diversification estimators and clade delimitations.** Plot elements follow Figure 3.

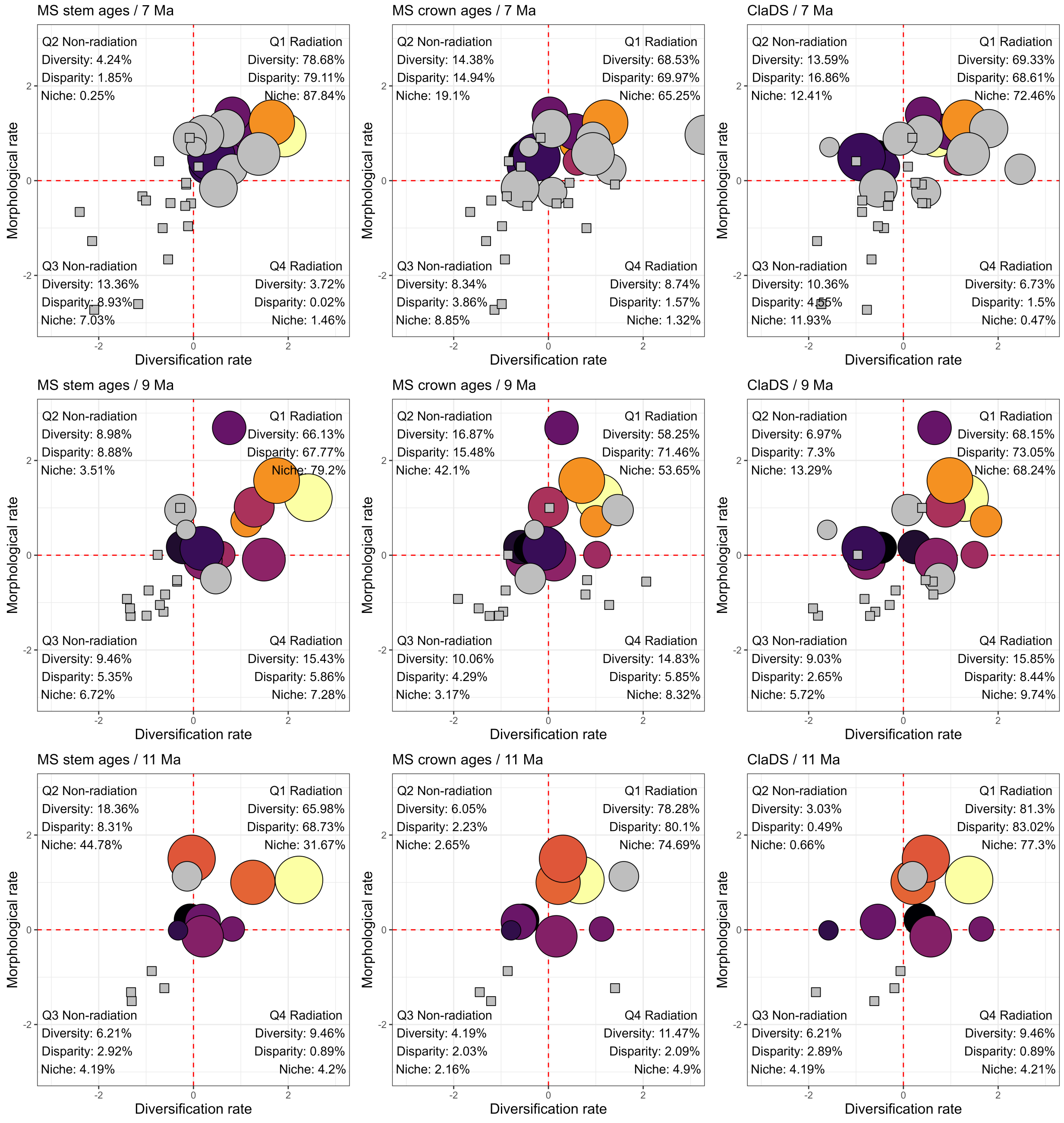

**Fig. S7: The radiation space of *Mimosa* under the 9 Ma clade delimitation.** (a) Plot showing clade disparity and niche breadth, along with biodiversity metrics for each quadrant. (b) Plot showing the clade numbers under the 9 Ma clade delimitation. Taxa belonging to each clade are depicted in Fig. S3.

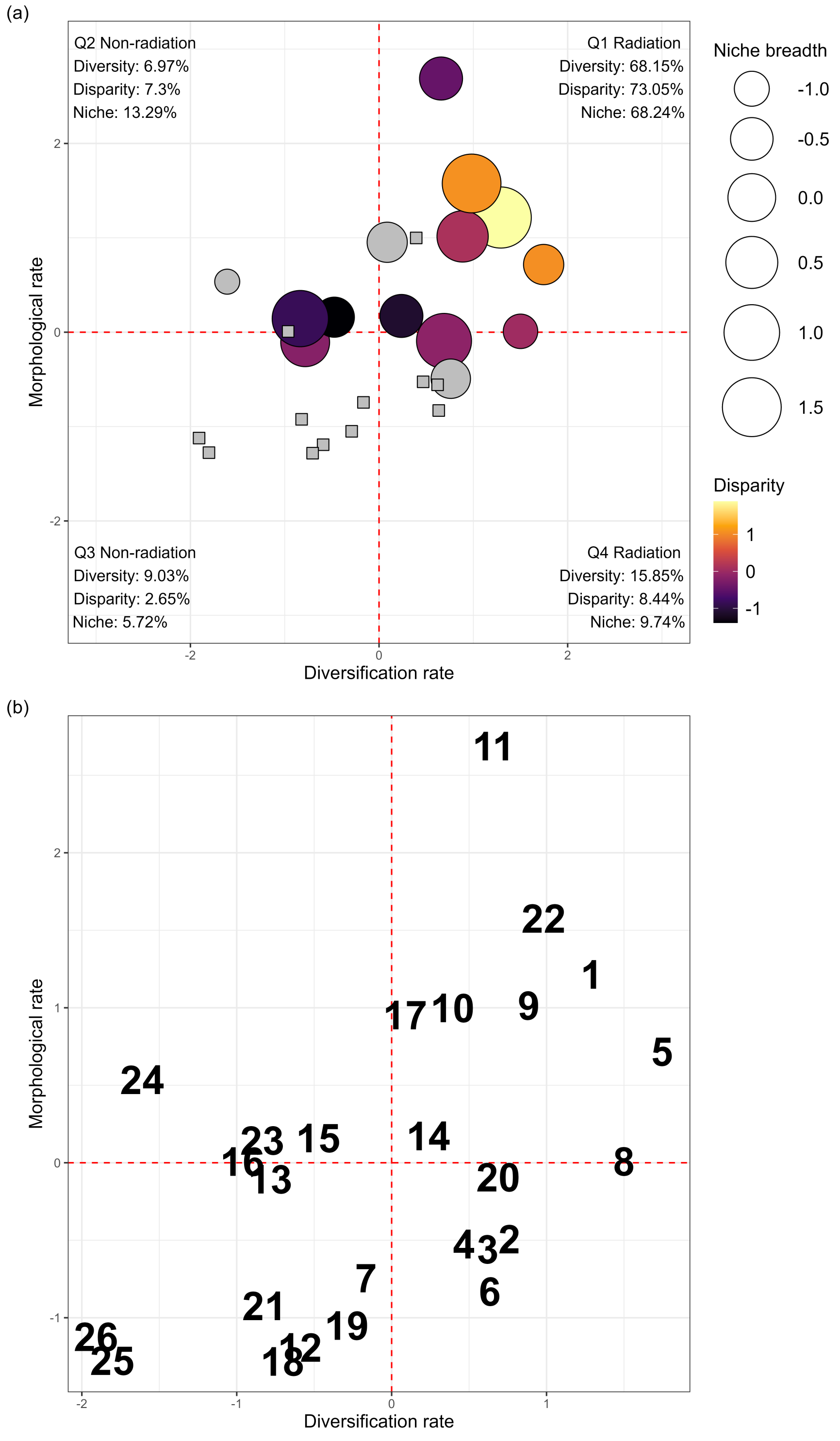

**Fig. S8: Rates of net diversification and altitudinal niche change across the *Mimosa* phylogeny under the 9 Ma clade delimitation.** Tip labels are shown to facilitate the identification of *Mimosa* clades from Simon et al. (2011) and Vasconcelos et al. (2020), as depicted in Fig. S3. Net diversification rates plotted here were estimated using the ClaDS model for the whole tree, then averaged within each clade. Both rates were mean-centered and scaled by their standard deviations.

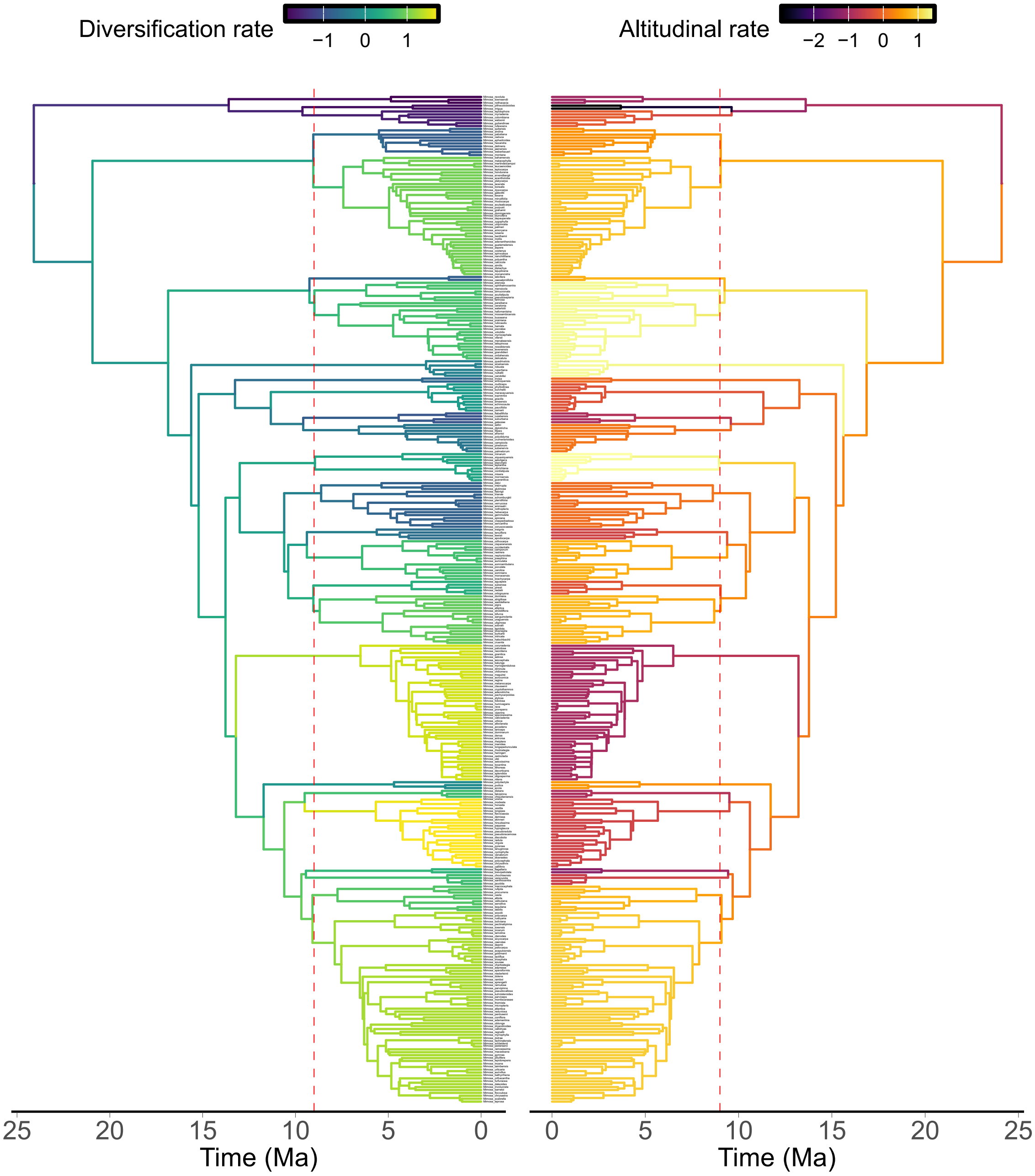

### Supplementary Tables

**Table S1: Vegetative morphological traits compiled from the literature.** Trait nomenclature and descriptions follow Barneby (1991).

| Trait | Description |
|---|---|
| Leaf-stalk length | Length of the entire primary leaf axis, from its insertion on the stem to its distal end. |
| Pinna rachis length | Length of the secondary leaf axis, from its insertion on the primary axis to its distal end. |
| Leaflet length | Length of the leaflet, from base to apex. |
| Leaflet width | Maximum width of the leaflet, measured perpendicular to its length. |
| Number of pinna pairs | Total number of opposite pinna pairs along the primary leaf axis. |
| Number of leaflet pairs per pinna | Total number of opposite leaflet pairs along each pinna rachis. |
| Plant height | Vertical distance from the ground level to the highest point of the plant. |

**Table S2: pGLS analyses of ClaDS diversification rates as a function of niche change rates across clade delimitations.** *P*-values are two-sided, based on Wald tests of the regression coefficients. **P* < 0.05.

| Model | Delimitation (Ma) | Slope | Adjusted $R^2$ | *P*-value |
|---|---|---|---|---|
| Diversification ~ Altitudinal rate | 7 | 0.390 | 0.170 | 0.008* |
| Diversification ~ Climatic rate | 7 | 0.151 | 0.006 | 0.281 |
| Diversification ~ Edaphic rate | 7 | 0.341 | 0.137 | 0.016* |
| Diversification ~ Altitudinal rate | 9 | 0.204 | 0.018 | 0.240 |
| Diversification ~ Climatic rate | 9 | -0.030 | -0.040 | 0.850 |
| Diversification ~ Edaphic rate | 9 | 0.287 | 0.086 | 0.080 |
| Diversification ~ Altitudinal rate | 11 | 0.043 | -0.088 | 0.872 |
| Diversification ~ Climatic rate | 11 | -0.147 | -0.051 | 0.531 |
| Diversification ~ Edaphic rate | 11 | 0.221 | -0.025 | 0.419 |

**Table S3: pGLS analyses of Magallón and Sanderson estimator diversification rates as a function of niche change rates across clade delimitations.** Diversification rates were estimated using crown or stem ages, and with an extinction value of 0.5. *P*-values are two-sided, based on Wald tests of the regression coefficients. **P* < 0.05.

| Model | Delimitation (Ma) | Slope | Adjusted $R^2$ | *P*-value |
|---|---|---|---|---|
| Diversification (crown) ~ Altitudinal rate | 7 | 0.373 | 0.113 | 0.027* |
| Diversification (crown) ~ Climatic rate | 7 | 0.207 | 0.017 | 0.217 |
| Diversification (crown) ~ Edaphic rate | 7 | 0.394 | 0.130 | 0.019* |
| Diversification (stem) ~ Altitudinal rate | 7 | 0.391 | 0.127 | 0.020* |
| Diversification (stem) ~ Climatic rate | 7 | 0.176 | 0.004 | 0.291 |
| Diversification (stem) ~ Edaphic rate | 7 | 0.404 | 0.138 | 0.016* |
| Diversification (crown) ~ Altitudinal rate | 9 | 0.034 | -0.041 | 0.872 |
| Diversification (crown) ~ Climatic rate | 9 | -0.170 | -0.010 | 0.396 |
| Diversification (crown) ~ Edaphic rate | 9 | 0.292 | 0.047 | 0.148 |
| Diversification (stem) ~ Altitudinal rate | 9 | 0.425 | 0.146 | 0.031* |
| Diversification (stem) ~ Climatic rate | 9 | 0.140 | -0.021 | 0.494 |
| Diversification (stem) ~ Edaphic rate | 9 | 0.431 | 0.152 | 0.028* |
| Diversification (crown) ~ Altitudinal rate | 11 | 0.196 | -0.049 | 0.520 |

**Table S3.** (Continued)

| Model | Delimitation (Ma) | Slope | Adjusted $R^2$ | *P*-value |
|---|---|---|---|---|
| Diversification (crown) ~ Climatic rate | 11 | -0.089 | -0.082 | 0.773 |
| Diversification (crown) ~ Edaphic rate | 11 | 0.546 | 0.234 | 0.054 |
| Diversification (stem) ~ Altitudinal rate | 11 | 0.180 | -0.055 | 0.556 |
| Diversification (stem) ~ Climatic rate | 11 | 0.034 | -0.090 | 0.912 |
| Diversification (stem) ~ Edaphic rate | 11 | 0.305 | 0.010 | 0.311 |

**Table S4: pGLS analyses of morphological evolution rates as a function of niche change rates across clade delimitations.** *P*-values are two-sided, based on Wald tests of the regression coefficients. **P* < 0.05.

| Model | Delimitation (Ma) | Slope | Adjusted $R^2$ | *P*-value |
|---|---|---|---|---|
| Morphological rate ~ Altitudinal rate | 7 | 0.452 | 0.180 | 0.006* |
| Morphological rate ~ Climatic rate | 7 | 0.413 | 0.145 | 0.014* |
| Morphological rate ~ Edaphic rate | 7 | 0.520 | 0.248 | 0.001* |
| Morphological rate ~ Altitudinal rate | 9 | 0.350 | 0.086 | 0.079 |
| Morphological rate ~ Climatic rate | 9 | 0.279 | 0.040 | 0.167 |
| Morphological rate ~ Edaphic rate | 9 | 0.348 | 0.085 | 0.081 |
| Morphological rate ~ Altitudinal rate | 11 | 0.150 | -0.067 | 0.626 |
| Morphological rate ~ Climatic rate | 11 | 0.151 | -0.066 | 0.622 |
| Morphological rate ~ Edaphic rate | 11 | 0.280 | -0.005 | 0.353 |
